# Representational invariance of numerosity via divisive normalization in coarse and fine visual channels

**DOI:** 10.64898/2026.02.21.707228

**Authors:** Joonkoo Park, Shimin Hu

**Author notes:** Corresponding author: Joonkoo Park, Ph.D., 135 Hicks Way, Amherst MA 01003.

## Abstract

Robust evidence now suggests that numerosity perception emerges from the early visual cortex. However, such empirical findings pose a theoretical challenge for explaining how a low-level perceptual system represents discrete values from continuous input independently of other magnitude dimensions. Among proposals for this representational invariance, two computational accounts with ties to neural data have garnered attention: divisive normalization and Fourier decomposition. Here, we test these hypotheses using an integrated neural, behavioral, and computational approach and show that: (1) The visual cortex remains sensitive to numerosity even when the input images are equalized for their Fourier power, inconsistent with the Fourier decomposition account. (2) The divisive normalization model explains this neural phenomenon through the selective disruption of fine but not coarse visual channels when encoding normalized local contrast. (3) Backward masking that disrupts fine processing in a psychophysical experiment degrades the acuity of intact dot arrays to the level of Fourier-power equalized dot arrays, which validates the unique prediction of the divisive normalization model. These findings provide converging evidence for the proposal that normalized local contrast enables representational invariance of numerosity in the visual cortex.

## Introduction

Numerosity perception has been a central topic of research in psychology, neuroscience, and philosophy due to the fundamental challenge of achieving representational invariance (Burr et al., 2018; Clarke & Beck, 2021; Nieder, 2025). How does the perceptual system represent the number of items in a collection irrespective of other confounding magnitude dimensions such as size of the items or the density of the collection? Understanding this perceptual mechanism is a significant challenge from a neurocomputational perspective because one must identify a mechanism that extracts discrete numerical values from continuous sensory measures while maintaining invariance to other dimensions.

A wealth of empirical research has now demonstrated that this representational invariance is achieved early in the visual system. For example, human imaging studies show that neural activity in early visual cortex (as early as V1) is sensitive to changes in numerosity of the stimulus with relatively little sensitivity to other dimensions (Park et al., 2016; Fornaciai et al., 2017; DeWind et al., 2019; Castaldi et al., 2019; Kido et al., 2025; Karami et al., 2025; Zhang et al., 2025). Consistent with these neural findings, psychophysical studies have shown that numerosity judgment is relatively independent of other magnitude dimensions and is susceptible to various perceptual effects, such as adaptation and serial dependence, that are thought to operate in early visual areas (Anobile et al., 2016; Cicchini et al., 2016; DeWind et al., 2015; Fornaciai & Park, 2018). Unlike this accumulating evidence in empirical studies, however, the mechanism underlying this representational invariance remains to be elucidated at the theoretical level.

Among various accounts developed in the past (Dakin et al., 2011; Dehaene & Changeux, 1993; Stoianov & Zorzi, 2012), two recent lines of work have proposed specific mechanisms with ties to neural data for how this representational invariance is achieved in the brain. The first proposal (Park & Huber, 2022) contends that divisive normalization between neurons with center-surround receptive fields (**Fig. 1A**) effectively normalizes neuronal activities across dimensions that are orthogonal to numerosity, thereby making the pool of neurons most sensitive to numerosity. Thus, according to this proposal, *normalized local contrasts* of the image make up the sensory representation of numerosity. The second proposal (Paul et al., 2022) posits that the early visual system approximates a Fourier decomposition (**Fig. 1B**) and argues that the summation of “the absolute power spectral density (PSD) of the Fourier transform” tracks numerosity with little effect of object size and spacing. Thus, according to this proposal, aggregate Fourier power of the image makes up the sensory representation of numerosity.

**Figure 1.**
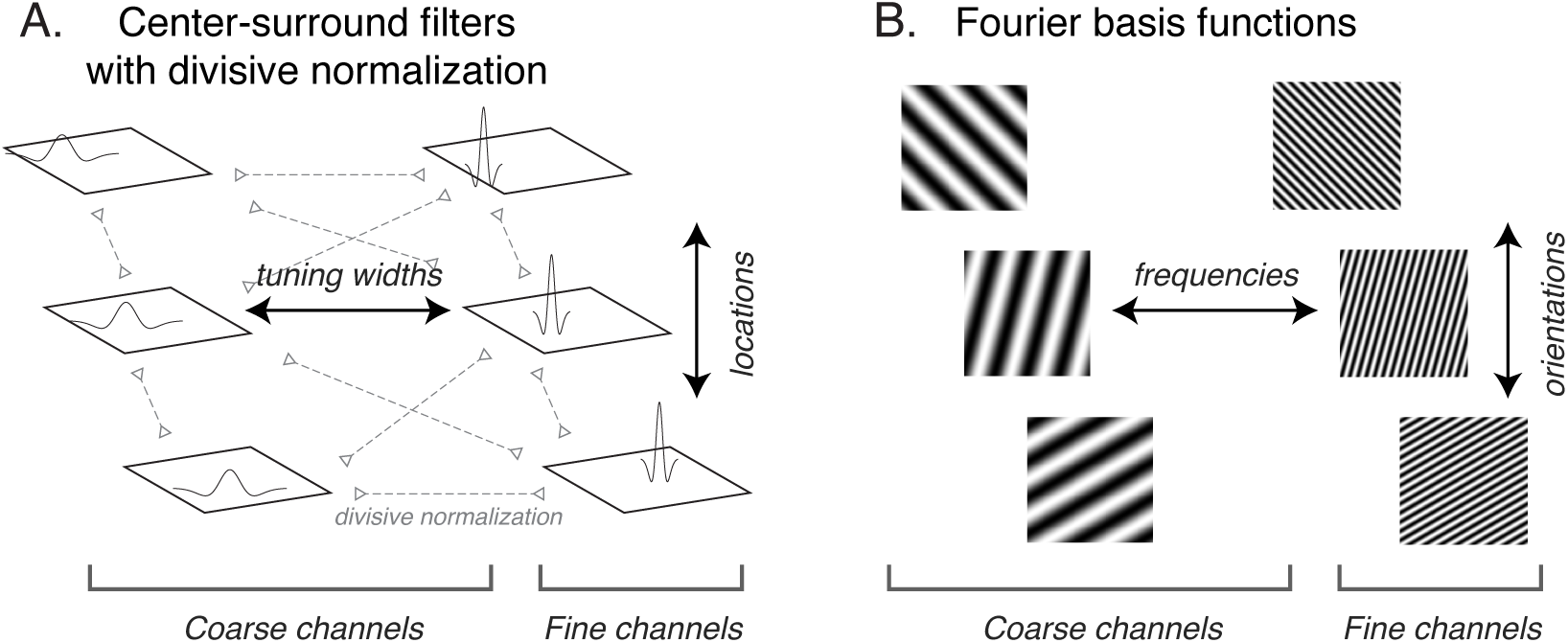
Schematic illustration of two hypothetical mechanisms for the sensory encoding of numerosity. A. In the divisive normalization model, input images are encoded by center-surround filters across the visual field at multiple spatial scales which are mutually inhibitory (Park & Huber, 2022). B. In the aggregate Fourier power model, input images are decomposed via Fourier analysis into basis functions across frequencies and orientations (Paul et al., 2022).

The current study aims to put these hypotheses to a rigorous test. In their previous work, Park & Huber (2022) criticized Paul and colleagues’ (2022) proposal on a theoretical basis. Namely, it was argued that the way aggregate Fourier power is computed for an image was flawed from a neurobiological point of view. However, these theoretical criticisms did not accompany empirical validations. In this study, using an integrated neural, behavioral, and computational approach, we provide an empirical test for the Fourier decomposition account. To do so, we first constructed a set of dot array images ranging in numerosity, size, and spacing (DeWind et al., 2015). Then, we artificially equated Fourier power across all those images as in a previous behavioral study (Adriano et al., 2021b; Willenbockel et al., 2010). We reasoned that if the initial sensory representation of numerosity is based on Fourier decomposition of the visual system, then the brain should be insensitive to numerosity of the dot array images whose aggregate Fourier power are equalized.

It is worth noting that the ‘dots’ in the Fourier-power-equated (FPE) images are still identifiable despite some visual noise (see **Fig. 2**). Indeed, previous behavioral studies showed that participants are nonetheless able to make numerosity judgments following Weber’s law with FPE images (Adriano et al., 2021a, 2021b). Such a result already suggests that aggregate Fourier power is unlikely to serve as a sensory representation of numerosity. Nevertheless, how the brain becomes sensitive to numerosity despite FPE remains to be understood.

**Figure 2.**
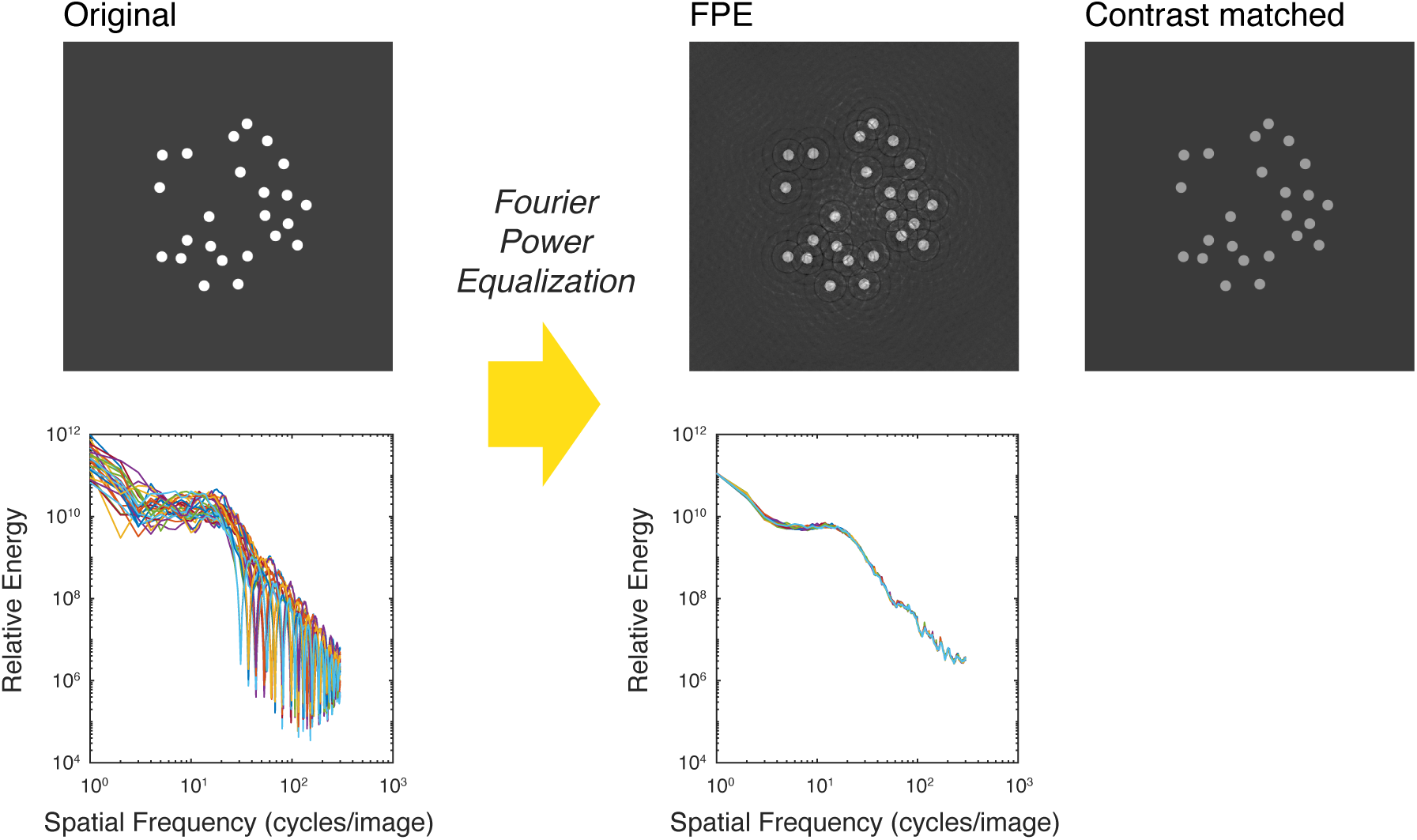
Stimulus construction with examples. The Original dot array images were submitted to Fourier power equalization the SHINE toolbox where the rotational average of the Fourier energy spectrum across all the images were matched. This procedure resulted in the Fourier power equalized (FPE) images. In the computational modeling and the psychophysical experiment part of the study, the foreground and background gray scale values of the Original images were adjusted to match their average gray scale values of the FPE images, which yielded the Contrast matched images.

Here, we first conduct a visual evoked potential (VEP) experiment to identify the temporal cascade of neural activities in response to intact and FPE images of dot arrays, in the absence of any magnitude-related judgment. The results suggest a strong disruption in the fine visual channels for the FPE images, thereby suggesting the need for distinguishing fine and coarse channels in numerosity perception. Following this insight, we adopt the divisive normalization model (Park & Huber, 2022) to propose a possible explanation for the VEP data in light of coarse and fine visual channels. This explanation leads to a novel prediction according to which experimental disruption of the fine channels should selectively reduce the sensitivity to numerosity at the behavioral level. We then test this novel prediction using a psychophysical experiment. Our results, as reported below, rule out the aggregate Fourier power account of numerosity perception and suggest normalized local contrasts (acquired via divisive normalization) across multiple spatial-scale channels as the key candidate for the initial sensory representation of numerosity.

## Materials and Methods

### Electroencephalography (EEG) experiment

#### Participants

Thirty-five participants were originally recruited from the departmental research participation system for course credits. Data from one subject were not recorded due to experimenter error, making the final sample size to be N=34. Among these participants, 6 identified as males and 28 as females. The age range was from 18.5 to 22.1 years with the mean of 20.1 years. Self-reported ethnic and racial category were as follows: Hispanic or Latino (4), Not Hispanic or Latino (28), No Response (2); White or Caucasian (29), Asian (5). All participants were right-handed, except for one who reported as ambidextrous. All participants were naive to the purpose of the experiment and had normal or corrected-to-normal vision. The experimental protocol was approved by the University of Massachusetts Amherst Institutional Review Board, and all participants provided written informed consent.

#### Stimuli

The experiment used two types of images: Intact images and Fourier amplitude spectra equalized images. These images were presented on a monitor screen running at 144 Hz with a resolution of 1920×1080, encompassing approximately 34°×19° of visual angle from the viewing distance of about 90 cm.

The Original dot-array images initially contained white dots (#FFFFFF) on a dark-gray background (#555555) of size 600×600 pixels. Following the dot-array generation scheme developed by DeWind and colleagues (2015), the images were systematically constructed to span three levels of number (*N*), size (*Sz*), and spacing (*Sp*) in a logarithmic scale, making it a total of 27 unique sets of parameters. Under this scheme, *N* ranged from 8 to 32 dots, the diameter of a dot ranged from 12 to 24 pixels (0.20° visual angle on average). The diameter of the field (within which the dots were drawn) ranged from 180 to 360 pixels (4.52° visual angle on average). The dot arrays were expected to project onto the parafovea. The area of each dot within an array was identical, and the minimum center-to-center distance between any two dots was at least the dot diameter. Thirty-two unique images were generated for each of the 27 parameter sets (making a total of 864 images). All the stimuli were generated using the published code for generating dot arrays for magnitude perception studies (https://osf.io/s7xer/).

The Fourier amplitude spectra equalized images (henceforth, Fourier power equalized or FPE images for simplicity) were created from the Original images using the SHINE toolbox (Willenbockel et al., 2010). This toolbox allows for the equalization of low-level visual attributes, such as luminance and Fourier spectra, across images. Specifically, we used this toolbox so that the rotational average of the Fourier amplitude spectra (*sfMatch*) and the luminance histogram (*histMatch*) of all 864 Intact images were matched. In the EEG experiment, the Original images were used in the Intact blocks of the experiment. Examples of the images used in this study are shown in **Figure 2**.

#### Procedure

The EEG experiment was presented using Psychtoolbox 3 on MATLAB (R2013a). The EEG data was continuously recorded using an active electrode amplifier (actiCHamp, Brain Products, GmbH) from 64 channels distributed in an extended coverage, triangulated, equidistance cap (M10, EasyCap, GmbH), with a sampling rate of 1000 Hz. Each channel was labeled according to its nearest neighbor in the standard 10-5 nomenclature. During the recording, all the channels were referenced to the vertex (Cz). One vertical EOG electrode was placed below the left eye and two EOG electrodes were placed lateral to the left and right canthi. Channel impedances were kept below 15 kΩ most of the time, although up to 35 kΩ was tolerated.

Each participant completed four Intact blocks and four FPE blocks; some participants received the Intact blocks first and the other half received the FPE blocks first, in a counterbalanced manner. Each block contained 378 trials. On each trial, the stimulus (Intact or FPE image) was presented for 200 ms followed by an intertrial interval (ITI) chosen from a uniform distribution ranging [500 700] ms. A black fixation cross was presented at the center of the screen throughout the trial. Participants were instructed to press a button under their right or left index finger (counterbalanced) when this fixation cross turned red, which happened in 27 pseudorandom trials within each block. The fixation cross turned yellow in response to participant’s button press. These oddball trials were excluded from the event-related potentials analysis. All the participants were reasonably accurate in detecting the fixation color change with the average hit rate of 93.2% with the average (± SD) of the median RT being 454 ± 47 ms, suggesting that the participants on average were paying attention to the screen. Each block took about 5 minutes and participants took self-paced breaks in between blocks.

#### Data analysis

The EEG data were analyzed off-line using EEGLAB and ERPLAB (Delorme & Makeig, 2004; Lopez-Calderon & Luck, 2014). The raw continuous EEG data were high pass filtered at 0.01 Hz and referenced to the average value of all the 64 channels. The continuous data were then segmented into epochs ranging [-100 500] ms from stimulus onset, with a pre-stimulus-interval baseline correction. The epochs from both the Intact and FPE blocks were combined prior to the independent component analysis (ICA) to identify artifactual (i.e., eye blinks) components. Then, the ICA weights representing the artifacts were removed from all the epochs for Intact and FPE blocks. Then, epochs with signal going outside the range of [-150 150] μV were further rejected using an ERLPAB artifact rejection tool. This procedure led to the median rejection rate of 5.4% in Intact blocks and 6.5% in FPE blocks. Finally, the epochs were selectively averaged for each of the 27 parameters sets (across three levels of *N*, *Sz*, and *Sp*, respectively), followed by a low-pass filter at 30 Hz.

These event-related potentials (ERPs) data at each latency point and at each channel were submitted to linear mixed-effects model analysis using the *fitlme* function in MATLAB. The full model consisted of mean centered log-transformed *N*, *Sz*, and *Sp* values as fixed-effects variables and participants as a random-effects variable. The marginal R^2^ of the full model (i.e., the variance explained by the fixed-effects variables) was computed following (Nakagawa & Schielzeth, 2013). The statistical significance of the fixed-effects variables combined was computed by the likelihood ratio test using the difference of the deviances between this full and the reduced model which contained only the participants variable. To be maximally conservative considering the multiple comparisons issue, the Benjamini-Hochberg procedure was applied to p-values across all the channels and latency points with a false discovery rate of *q_FDR_* < 0.001, and statistical significance was considered only when this threshold was met for two or more consecutive latency points.

### Computational modeling

The current study adopted the previously published divisive normalization model for numerosity perception (Croteau et al., 2024; Park & Huber, 2022). Because the mathematical basis of the model has been elaborated in prior publications, here we primarily focus on the conceptual explanation of the model. Unless otherwise noted, the same parameters from Park & Huber (2022) were used in the current study. For more details, please refer to Park & Huber (2022).

#### Stimuli

The goal of this modeling work was to simulate the encoding of Intact and FPE images as used in our EEG experiment. To that end, we first prepared the dot-array stimuli used in our previous modeling work (Park & Huber, 2022) as the Original images. These images contained white dots on a black background (200×200 pixels) that spanned systematically across five-levels of *N*, *Sz*, and *Sp* dimensions, making it a total of 35 unique sets of parameters. *N* ranged from 5 to 20 dots, the diameter of a dot ranged from 9 to 18 pixels, and the field radius ranged from 45 to 90 pixels. Twenty unique dot-array images were created for each of the 35 parameter sets. These Original images underwent the aforementioned procedure in the SHINE toolbox to equalize their Fourier amplitude spectra and luminance histogram, which yielded the FPE images. This process decreased the overall contrast of the FPE images compared to the Intact images. In the computational stimulations and in the behavioral experiment, we intended to match the overall contrast between the two types of images. This was because the images were presented in an intermixed fashion, and we did not want differences in the results to be attributed to the overall contrast differences. Thus, the Original images were further adjusted so that the average gray scale levels of their foreground (dots) and background were matched to those of FPE images. Those contrast-adjusted images were used as Intact images.

#### Convolution and divisive normalization

As in our original neural network model (Park & Huber, 2022), the current architecture consisted of a set of six convolutional layers with varying sizes of difference-of-Gaussians (DoG) filters. The size of the filters was determined so that the smallest filters can pick out fine edges of the dots and the largest filters can pick out the overall landscape of the dots. Following the original work, the filter sizes were set to σ = {1, 2, 4, 8, 16, 32}, where σ is the standard deviations of the narrower Gaussian of the DoG. In our implementation, these filter sizes empirically resulted in the full zero-crossing width of approximately 3.4, 6.9, 13.8, 27.9, 56.1, and 112.1 pixels. Given these widths, the two narrowest filters would capture edges of a typical dot, and the rest of the filters would cover a whole dot or multiple dots and thus capture blobs of the dot array image. The largest filter would cover approximate half of the field area of the dot array. The output of DoG convolution underwent half-wave rectification, which was interpreted as the driving input, *D_i,k_*, for each neuron, *i* (∈{1, 2, … 200×200}), in filter size *k* (∈{1, 2, …, 6}).

At first, we simulated the encoding of Intact and FPE images using the original model where divisive normalization occurs simultaneously across all filter sizes. In this case, the driving input in each neuron, after some amplification, was normalized by the summation of driving inputs of neighboring neurons, *j*, weighted according to their Euclidean distances. That is, the normalized response was computed as follows:

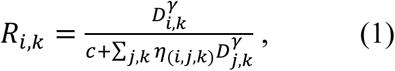

where 𝛾 (=2) indicates the amplification factor, *η* indicates the exponentially decaying neighborhood weight function, and *c* is a constant set as 1. Then, 𝛴*_i,k_ R* represented the aggregate magnitude of the simulated neuronal activity in response to the given input image.

#### Separating coarse and fine channels

After using our original formulation where divisive normalization was applied simultaneously across all filter sizes (Park & Huber, 2022), we next explored options to simulate coarse and fine channels separately. It is now well demonstrated in both single-cell studies and neuroimaging studies that the visual system processes coarse (low spatial frequency) and fine (high spatial frequency) information independently in parallel channels (Hegdé, 2008; Hochstein & Ahissar, 2002). To this end, we simulated the divisive normalization model separately for the coarse channels that encoded ‘blobs,’ and fine channels that encoded ‘edges’ (Schyns & Oliva, 1994). The coarse channels included DoG filters (more specifically the full zero-crossing width) that were larger than the smallest dot diameter (9 pixels) which led to *k* ∈{3, 4, 5, 6}, and the fine channels included the DoG filters that were smaller than the smallest dot diameter which led to *k* ∈{1, 2}.

### Psychophysical experiment

#### Participants

One hundred participants (N = 100; 22 identified as males and 78 females) participated in exchange for course credit (age range from 21 to 26 years with the mean of 24 years). Self-reported ethnic and racial category were as follows: Hispanic or Latino (9), Not Hispanic or Latino (91); White or Caucasian (61), Asian (24), Black or African American (10), Other (5). All participants were naive to the purpose of the experiment and had normal or corrected-to-normal vision. The experimental protocol was approved by the University of Massachusetts Amherst Institutional Review Board, and all participants provided written informed consent.

#### Stimuli

Original dot-array images with white dots (#FFFFFF) on a dark-gray background (#404040) of size 600×600 pixels were first created systematically across number (*N*), size (*Sz*), and spacing (*Sp*) (DeWind et al., 2015), as in the EEG experiment. This time, *N* ranged from 12 to 48 dots, the diameter of a dot ranged from 14 to 28 pixels, and the radius of the field (within which the dots were drawn) ranged from 120 to 240 pixels, in seven logarithmically equidistant levels. The Original dot arrays with median values of *N*, *Sz*, and *Sp* were used as the ‘reference’ stimuli for the 2AFC (two-alternative forced-choice) task (see *Procedure* below). Sixty-four unique reference images were generated. As in the EEG experiment, the FPE images were created from those Original images using *sfMatch* and *histMatch* in the SHINE toolbox. Then Intact images were created by matching the grayscale levels of the foreground and background of the Original images with the FPE images. The Intact and FPE images were used as the ‘test’ stimuli for the 2AFC task. Four unique test images were generated for each of the 91 parameter sets (making a total of 364 images).

#### Procedure

The psychophysical experiment was developed and presented using PsychoPy (version 2022.1.1). Each participant completed trials under 2×2 experimental conditions that differed in the test stimuli: (1) Intact images presented for 500 ms, (2) Intact images presented for 50 ms, (3) Fourier Power Equalized (FPE) images presented for 500 ms, and (4) FPE images presented for 50 ms. In all conditions, participants compared these test image to a fixed reference image with 24 dots. Specifically, each trial followed a fixed sequence: an Original reference image displayed for 250 ms, followed by a 100 ms mask; an interstimulus interval of 250 ms; a test image (either Intact or FPE) displayed for either 50 ms or 500 ms, followed by another 100 ms mask. After this sequence, participants were asked to press “1” to indicate the first image contained more dots or “2” to indicate the second image contained more dots. The intertrial interval, between the key press and the following trial, was a duration randomly chosen between 1400 ms and 1590 ms. There was a total of 8 blocks, each containing 89 trials. The backward visual masks were generated in PsychoPy using the *Noise* component. The noise type was binary, with nearest-neighbor interpolation and a noise element size of 10 px, producing a pixelated black-and-white square noise pattern. The mask was presented at a size of 600 × 600 pixels, and a central fixation cross was overlaid on the mask. These masks were designed to disrupt slower and more sustained visual channels of the target images, specifically in the 50 ms conditions (Breitmeyer & Ganz, 1976).

#### Data analysis

Behavioral responses were analyzed by fitting a cumulative Gaussian psychometric function separately for each experimental condition. The model included the log-transformed numerosity as the independent variable and individual participant’s choice response as the dependent variable, and it was implemented using the *quickpsy* toolbox in R (version 4.5.0), with the lapse rate set to 0.02 and all other parameters set to their default values (Linares & López-Moliner, 2015). From these models we obtained the point of subjective equality (PSE) and the just noticeable difference (JND) for each experimental condition for each participant. PSE refers to the *logN* value at which the observer is equally likely to choose either response. JND was calculated as the difference between the 75% threshold, where the observer is expected to be correct 75% of the time, and the PSE. Outliers were identified using individual Weber fraction (w). In each subject, w was computed by dividing JND by PSE for each experimental condition. Then, if w from any of the four experimental conditions exceeded the third quartile plus one-point-five times the interquartile range, the entire data from that subject was treated as outliers. Fifteen participants were removed from further analysis using this criterion. Individual estimates of PSE and JND were submitted to linear mixed-effects models using the *lmer* function from the *lmerTest* package in R. For visualization, we present the psychometric curves from the same model with the data across all participants collapsed.

## Results

### Qualitatively different ERP patterns for Intact and FPE images

We first present a gross report of the effects of *N*, *Sz*, and *Sp* of Intact and FPE images on the visual-evoked potentials. **Figure 3** illustrates the marginal R^2^, or the variance of the neural activities explained by *N*, *Sz*, and *Sp*, across all the channels. The Intact images elicited two distinct latency windows with significant R^2^ effects: one window centered around 100 ms and another window encompassing 200-300 ms. The earliest statistically significant latency point was at 68 ms in channel Oz and at 70 ms in channel PPO8h. The FPE images elicited a different pattern in which significant latency windows were extended across a much longer range from 150 to 450 ms. Statistically significant R^2^ effects were also found around 100 ms in the FPE images with the onset of statistical significance at 80 ms in the medial parietal channels centered on channel Pz; however, the effects at these early latencies were smaller than the case of Intact images. These results suggest that the brain represents the two types of images quite differently across the time course.

**Figure 3.**
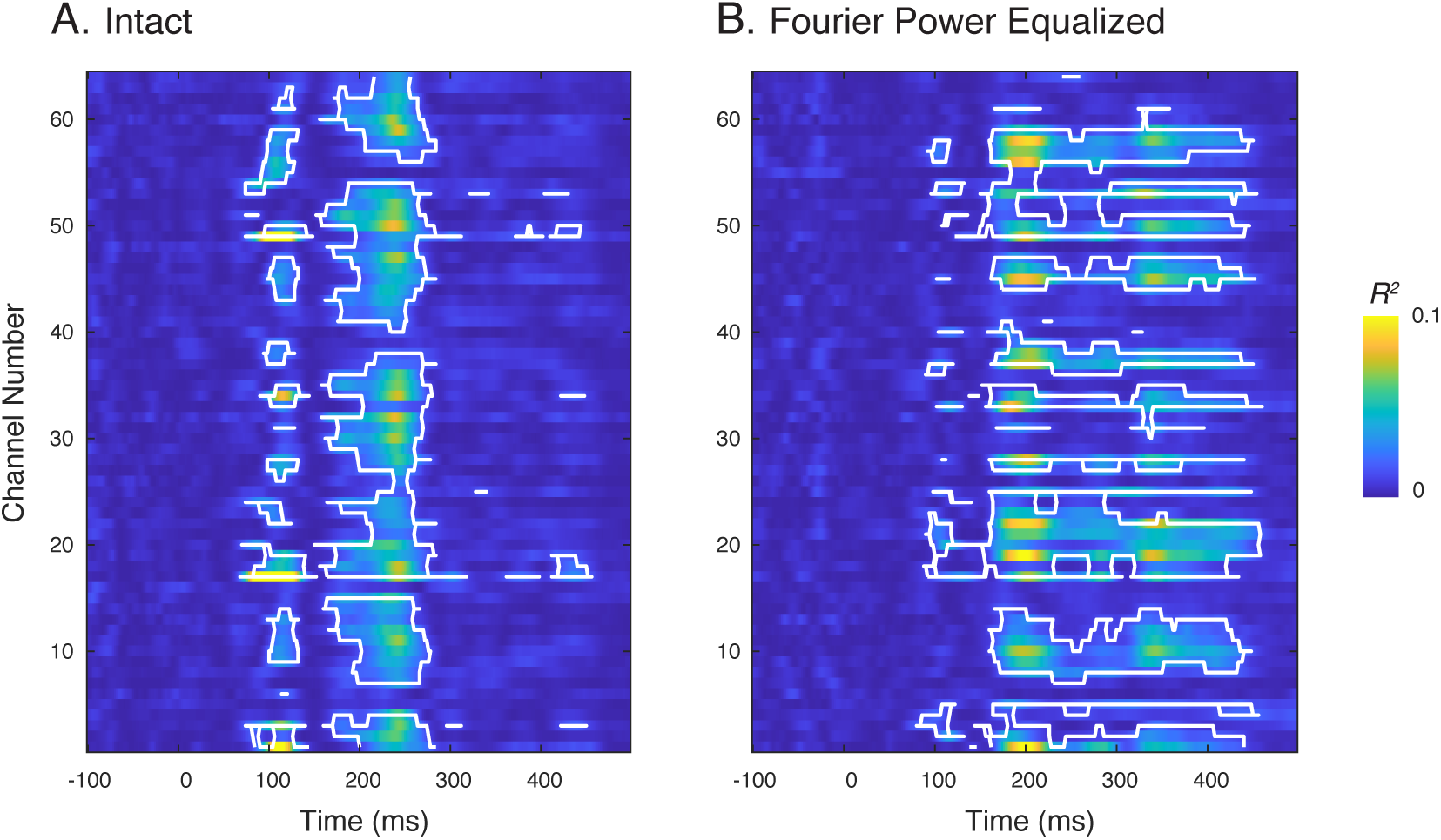
Gross graphical display of marginal R^2^ showing the variance of ERPs explained by the combined effects of N, Sz, and Sp from Intact (A) and FPE (B) images. White contour lines indicate statistically significant latency points under a conservative threshold (q_FDR_ < 0.001).

Next, we report topographic maps that offer a more detailed pattern of results (**Fig. 4**). In Intact images, early peaks of marginal R^2^ were driven by a strong effect of *N* and some effect of *Sp* in the medial occipital channels from 75-125 ms (mean values of beta estimates at channel 17 were *β_N_* = -1.3181, *β_Sz_* = -0.1660, *β_Sp_* = 0.3237). Later peaks of marginal R^2^ were driven by a similarly strong effect of *N* with some effect of *Sz* in slightly bilateral occipital channels from 175-275 ms (mean values of beta estimates at channel 50 were *β_N_* = 0.7916, *β_Sz_* = 0.3567, *β_Sp_* = -0.1569). These relatively strong effects of *N* in the two spatiotemporal locations replicate previous ERP findings of numerosity perception (Fornaciai et al., 2017; Park et al., 2016). In FPE images, the effect of *N* dominated throughout the time course as well. However, the direction, latency, and topography of the effects were different from the case of Intact images. First, although marginal R^2^ increased in the medial parietal channels around 75 ms (*β_N_* = 0.3453, *β_Sz_* = -0.0339, *β_Sp_* = 0.2425 at channel 17) and in the bilateral occipital channels around 100-125 ms (*β_N_* = -0.7224, *β_Sz_* = -0.0347, *β_Sp_* = -0.1574 at channel 20), the stronger effect of *N* in the occipital channels did not arise until 125 ms (*β_N_* = 0.6855, *β_Sz_* = -0.0391, *β_Sp_* = 0.4048 at channel 17). Then, the effect of *N* remained strong broadly across 175-450 ms in the inferior occipital channels and in the central channels (part of which could be a result of volume conduction). These results again suggest qualitatively different processing streams across the two types of images.

**Figure 4.**
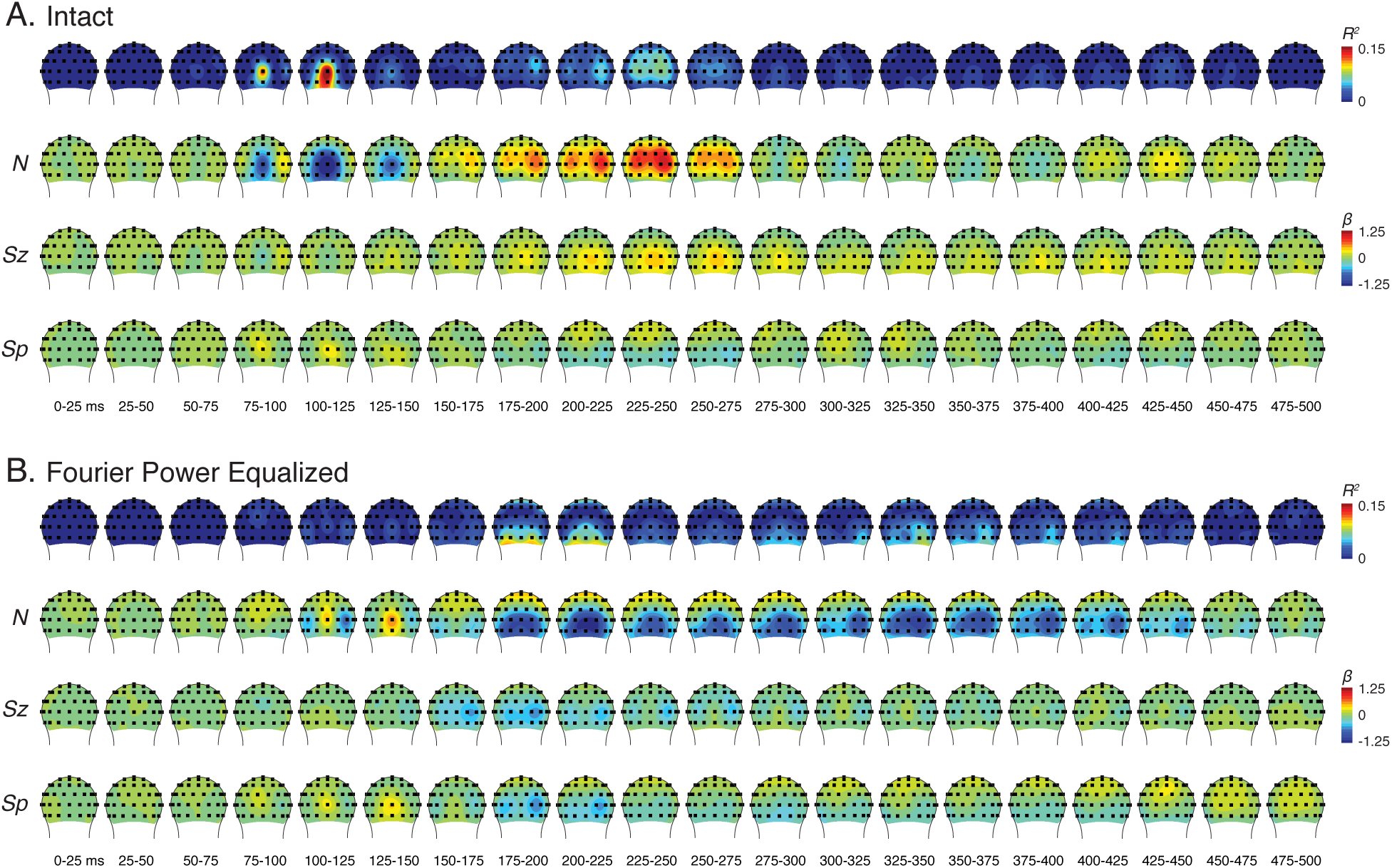
Posterior-view topographic maps of marginal R2, and the effect of N, Sz, and Sp on the ERPs from Intact (A) and FPE (B) images.

Finally, we assessed raw waveforms in response to the Intact and FPE images. Considering the dominant effects of *N* across latencies, we focused on the grand averaged ERPs along low, medium, and high levels of numerosity (8, 16, and 32 dots) in the posterior channels. As **Figure 5** illustrates, the waveforms from the Intact and FPE images were quite different. The Intact images evoked a prototypical pattern (Fornaciai et al., 2017; Park et al., 2016) including the modulation of the C1 component by *N* in channel Oz and the P2 component in the bilateral (but more so in the right) occipital channels. The ERPs in response to the FPE images did reveal early medial occipital negativity (as in the C1 component); however, the shape of the waveforms was much distorted compared to the pattern observed in Intact images. In addition, the modulatory effect of *N* on C1 was much weaker in FPE images than in Intact images, confirming what is reported in **Fig. 2** and **Fig. 3**. Another pronounced difference between the two conditions was the robust effect of *N* across extended regions and latencies (200-500 ms) with reversed direction of the effect in FPE images.

**Figure 5.**
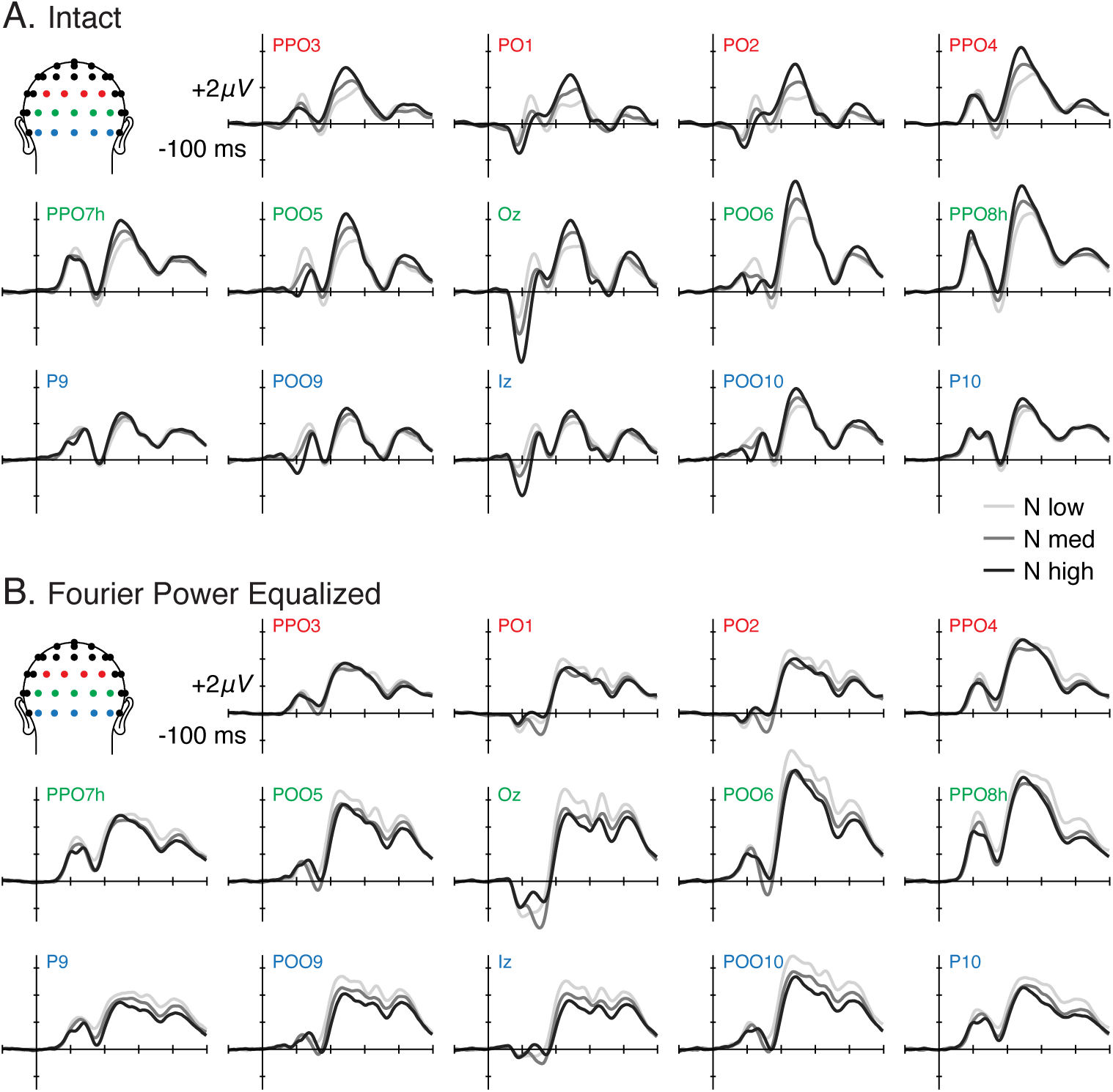
Posterior-channel waveforms of grand-averaged ERPs from Intact (A) and FPE (B) images as a function of low, medium, and high N. Channel labels are color coded to illustrate their topographical locations on the scalp.

Collectively, results from the EEG experiment demonstrated that Intact and FPE images yield qualitatively different neural activity patterns. The most prominent finding was the attenuation or absence of magnitude-related modulation of magnitude-related modulation (mostly by *N*) in the medial occipital C1 component for FPE images.

### Divisive normalization model simulates lack of numerosity modulation in FPE images

Our divisive normalization model (Croteau et al., 2024; Park & Huber, 2022) has been successful in simulating neural and behavioral data regarding the perception of numerosity of dot and item arrays. We first questioned what this original model would predict when given the FPE images. As shown in **Figure 6**, our model simulated a strong sensitivity to *N* and much weaker sensitivity to *Sz* and *Sp* of Intact images, replicating prior work. The slope of the linear fit to *logN*, *logSz*, and *logSp* were 10.823, 1.531, and 0.814, respectively. The FPE images, in contrast, yielded a different pattern. In this case, the slope of the linear fit to *logN*, *logSz*, and *logSp* were 0.777, -0.447, 0.122, respectively. While the relative insensitivity to *Sz* and *Sp* was consistent with the case of Intact images, the modulation by *N* was much reduced if any, with no clear monotonic response pattern as a function of *N*. The slope of the linear fit to *logN* in the Intact images (10.823) meant that the aggregate normalized response from Intact images increased by more than 10 times when *N* doubled; compared to this, the aggregate normalized response from FPE images changed barely as a function of *N*, *Sz*, or *Sp*. Another noticeable difference was the overall greater normalized responses compared to the case of Intact images, consistent with the greater overall ERP magnitudes in FPE images.

**Figure 6.**
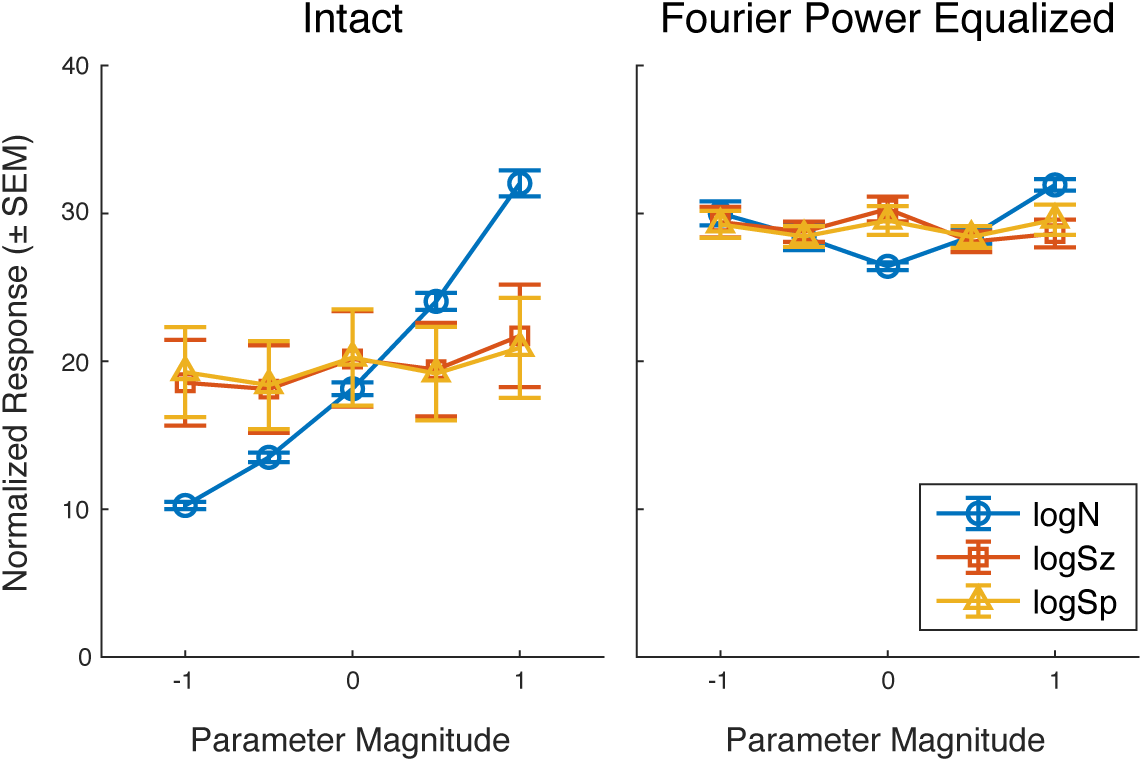
Aggregate normalized response as a function of parameters, N, Sz, and Sp (log2-transformed). The value 0 in the abscissa indicates median values of each dimension; 1 means twice of the median value; -1 means half of the median value. Error bars indicate standard error of the mean (SEM).

While the divisive normalization model correctly simulated the disruption of modulation by *N* at the level of C1 component, important questions remain. What aspects of the FPE images make such a drastic difference in both the computational model and the C1 activities, and what makes the subsequent neural activities (from 175 ms and on) still be modulated by *N*?

### Intact and FPE image processing differs in fine (but not in coarse) channels

Although Fourier power equalization did not change the gross profile of the amplitude spectra, it seriously disrupted the structure of the high (but not low) spatial frequency properties in each image (see **Fig. 2**). As a result, the FPE images lost sharp edges, leading to a reduction in local contrast and the prevalence of blurry textures throughout each image. Previous studies have shown that neural activities at the C1 level are strongly influenced by the parvocellular pathway, which is responsible for processing crips boundaries and strong local contrasts (Baseler & Sutter, 1997; Foxe et al., 2008; Hansen et al., 2011; Tobimatsu et al., 1995). Thus, our ERP results—particularly, the lack of C1 modulation by *N*—make sense in the context of the previous work. Similarly, our model simulations for the FPE images showed no effect of *N*, which would be consistent with the lack of C1 modulation by *N*. However, the ERP activities eventually turned out to be sensitive to *N*, yet the current model simulations do not explain that pattern. This invites a new way of thinking about modeling numerosity perception.

Decades of research suggests that the visual system processes low spatial frequency information (coarse features) and high spatial frequency information (fine features) in parallel channels with the coarse channel being processed earlier than the fine channel (Hochstein & Ahissar, 2002). Considering this prior literature, we explored one way to simulate coarse information processing separately from fine information processing. Specifically, we divided the divisive normalization model into two: the first, coarse model included filters whose full zero-crossing widths were larger than dot diameters thereby effectively representing blobs; the second, fine model included filters whose full zero-crossing widths were smaller than dot diameters thereby effectively representing edges (**Fig. 7**).

**Figure 7.**
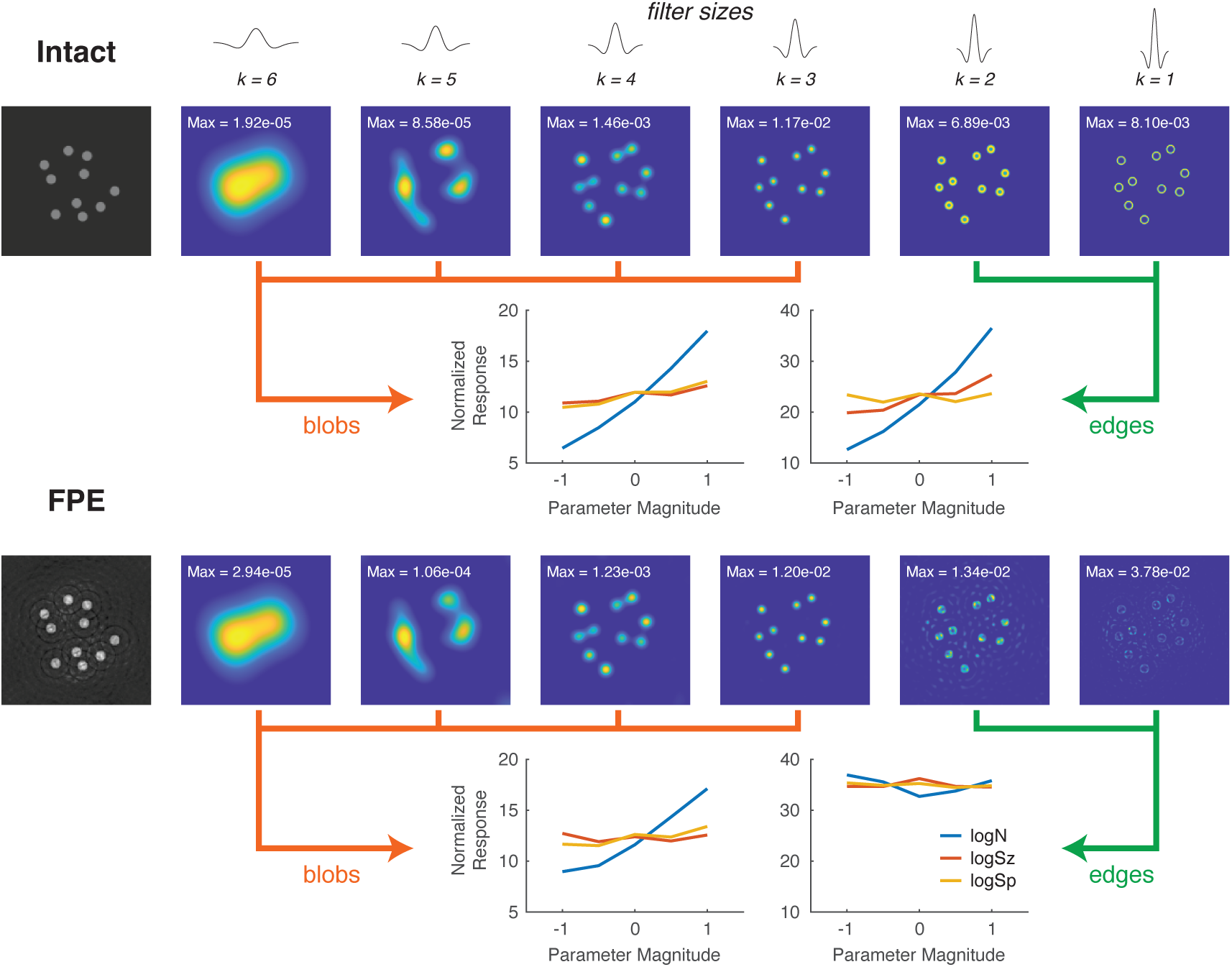
Divisive normalization model and the simulation results separately for coarse and fine filters. The maximum value labeled on each image represents the peak activation value in each layer and is reported to facilitate comparisons of relative responsivity across layers.

For the Intact images, both the coarse and the fine models showed a similar pattern where the normalized responses were modulated primarily by *N*. In the coarse model, the linear fit slopes for *logN*, *logSz*, and for *logSp* were 5.768, 0.802, and 1.259, respectively. In the fine model, the linear fit slopes for *logN*, *logSz*, and for *logSp* were 11.874, 3.638, and 0.111, respectively (**Fig. 7**). Note that one unit change in log-transformed total perimeter (TP) is equivalent to three quarters of a unit change in *logN* and one quarter of a unit change in *logSz*: *logTP* = log(2√𝜋) + ¼ × *logSz* + ¾ × *logN* (for derivation, see Park et al., 2016). Considering this relationship, normalized responses in the fine model closely matched total perimeter, which is not surprising given that the fine filters are exclusively sensitive to the edges.

In contrast to the Intact images, the FPE images yielded a different result. In the coarse model, the linear fit slopes for *logN*, *logSz*, and for *logSp* were 4.215, -0.050, and 0.866, respectively. In the fine model, the linear fit slopes for *logN*, *logSz*, and for *logSp* were -0.808, -0.051, and - 0.299, respectively (**Fig. 7**). These results suggest that, if visual processing of dot arrays is done through two parallel channels (as in our simulation), the coarse channel is sensitive to numerosity and related magnitudes like total perimeter in both Intact and FPE images, but the fine channel is sensitive to numerosity only in Intact images.

### Behavioral results match model predictions

Simulations from our computational model suggest one possibility in which both coarse and fine channels are modulated by *N* in the Intact images while only the coarse channel is modulated by *N* in the FPE images. This simulation outcome makes a couple of predictions. First, it predicts that the visual system is more sensitive to *N* when encoding Intact images than when encoding FPE images. In this scenario, observers making numerosity judgments would be more accurate with Intact images than with FPE images. Such a prediction had already been demonstrated, with more precise Weber fraction for Intact images than for FPE images (Adriano et al., 2021b). Our simulation makes another, novel prediction. If the processing of fine channels is disrupted (due to a psychophysical manipulation), the observer’s sensitivity to *N* will be selectively reduced for Intact images but not for FPE images. This is because the fine channels for the FPE images already do not show any sensitivity to *N*.

We tested this novel prediction using a backward masking paradigm. It is well accepted that visual stimuli evoke coarse channels much faster than they evoke fine channels (Maunsell & Gibson, 1992; Merigan & Maunsell, 1993; Shapley, 1990). According to influential models of visual masking (Breitmeyer & Ganz, 1976; Breitmeyer & Ogmen, 2000), the rapid and transient processing of the coarse features of a backward mask can disrupt the processing of slow and sustained processing of the fine features of the target stimulus under short stimulus onset asynchrony. In the psychophysical experiment, we used backward masking by noise to disrupt the processing of fine channels in Intact and FPE images. Participants judged which of the two arrays (‘reference’ followed by ‘probe’) contained more dots. The reference arrays were clear images with no Fourier power manipulations and always contained 24 dots. The probe arrays were either Intact or FPE images with varying *N*, *Sz*, and *Sp*. Importantly, in half of the trials the probe image was displayed for a very short duration (50 ms) immediately followed by a strong visual mask, making a conscious perception of the probe array difficult. In the other half of the trials, the probe image was displayed for a longer duration (500 ms); therefore, the probe array was clearly visible even though the same mask appeared afterwards. After all, it was a 2×2 design with image type (Intact vs. FPE) and image visibility (visible vs. masked) as factors.

The psychometric curve of the data collapsed across all the participants (**Fig. 8A**) show a ratio-dependent performance across all the conditions. Linear mixed-effects models were run on PSE and JND with image type and image visibility as fixed-effect regressors and participants as a random-effects variable. The linear model on PSE showed no significant effect of image type (F(1, 252) = 2.4497, p = 0.11880), image visibility (F(1, 252) = 3.2820, p = 0.07124), and the interaction (F(1, 252) = 3.1888, p = 0.07535). The linear model on JND showed a significant interaction effect (β = 0.03872, t(252) = 2.599, p = 0.009912). A follow-up computation of effect sizes using estimated marginal means (using *emmeans* in R) showed that for Intact images, visible dot arrays decreased JND relative to masked dot arrays (Cohen’s d = 0.36, 95% CI [0.07, 0.65]), while no reliable effect was observed for FPE images (d = 0.04, 95% CI [-0.33, 0.26]) (**Fig. 8B**).

**Figure 8.**
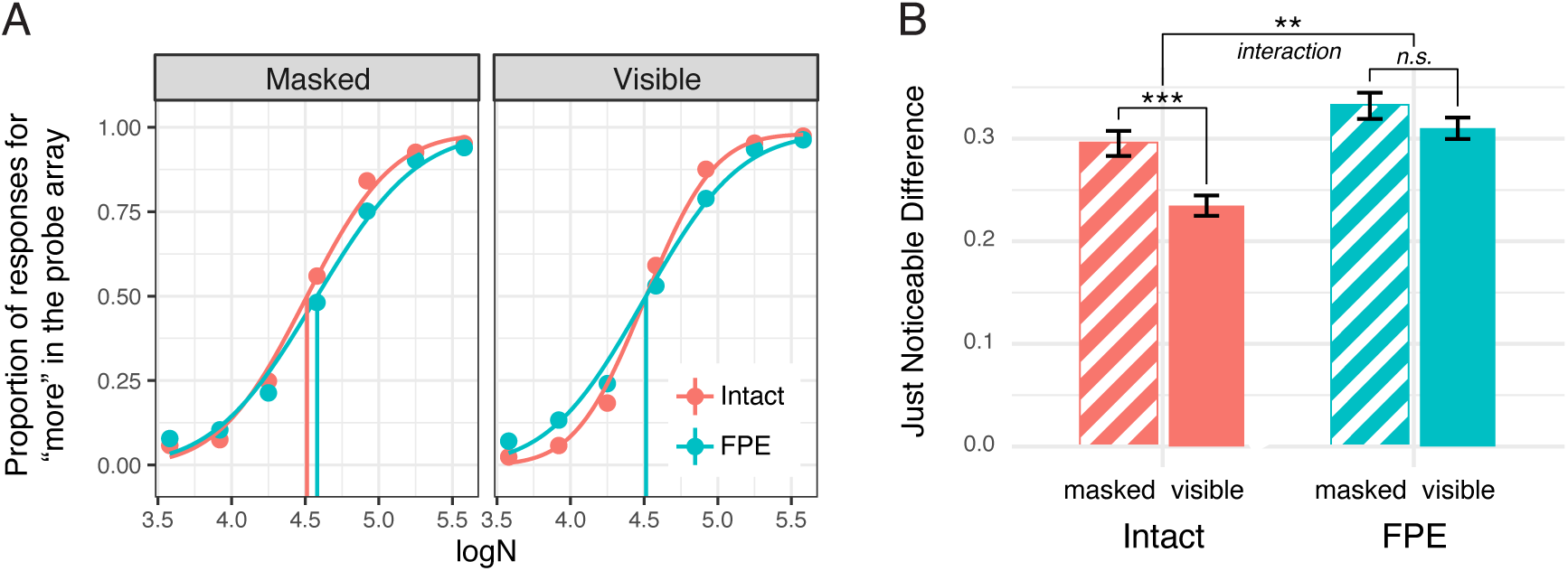
Results of the psychophysical experiment. A. Psychometric curves for numerosity comparison from data collapsed across all the participants. B. Average JND’s with standard error of the mean across four conditions. Smaller JND means greater sensitivity to numerosity. n.s., not significant; **, p < 0.01; ***, p < 0.001.

Post hoc t-tests of the various contrasts showed the following. Under the visible condition, the JND for the Intact images was significantly smaller (hence more sensitive to *N*) than the JND for the FPE images (β = 0.07544, t(84) = 7.095, p < 0.001). Under the masked condition, the JND for the Intact images was also significantly smaller than the JND for the FPE images (β = 0.03672, t(84) = 3.345, p = 0.001). For the Intact images, the JND was significantly smaller when the dot arrays were visible than when they were masked (β = -0.060591, t(84) = -6.662, p < 0.001). However, the JNDs were not significantly different for the FPE images (β = -0.02187, t(84) = -1.972, p = 0.052). These results from the JND analysis are consistent with the novel prediction developed from our computational model. When the processing of fine channels is disrupted by backward masking, participants’ sensitivity to *N* dropped substantially in the Intact condition but not in the FPE condition.

## Discussion

The current study was motivated to test two outstanding hypotheses regarding the sensory representational invariance of numerosity: the normalized local contrast account and the aggregate Fourier account. We first tested if and how Fourier-power-equalized (FPE) images modulated early visual cortical activities and found that the brain is still sensitive to the numerosity (and size and spacing to a smaller degree) of FPE images. The visual-evoked potential patterns suggested that FPE images disrupt information processing in the fine channels but not in the coarse channels, and this discrepancy could be explained by the divisive normalization model simulating coarse and fine channels separately. Our computational model together with the neural data suggested a coarse-to-fine processing in numerosity perception. Finally, the psychophysical study using backward masking confirmed the prediction made by the computational model concerning coarse-to-fine processing, that selective disruption of fine channels impedes participants’ sensitivity to intact images but not to FPE images because the information through the fine channels is indeterminant of numerosity in those images anyway. These findings have several implications. First, numerosity perception does not depend on aggregate Fourier power as suggested in a previous work (Paul et al., 2022). Second, numerosity perception like other visual features goes through coarse-to-fine processing stages. Finally, the predictive power of our computational model makes it one of the most powerful models that provide a mechanistic explanation of numerosity perception.

### Early visual activity for numerosity perception does not depend on aggregate Fourier power

There have been several computational accounts of achieving representational invariance of numerosity. In one of the earliest accounts, Dehaene and Changeux (Dehaene & Changeux, 1993) proposed that each object in a visual scene is encoded by a single neuron via a winner-take-all mechanism. This process effectively makes the neural network invariant to object location and size and thus achieves representational invariance of number. Dakin and colleagues (Dakin et al., 2011) proposed that the encoding of number is based on density perception which relies on the ratio between high spatial-frequency center-surround filter responses and low spatial-frequency center-surround filter responses. Stoianov and Zorzi (Stoianov & Zorzi, 2012) developed a hierarchical generative neural network model and showed that the activity from center-surround neurons receiving inhibition from neurons that encoded cumulative surface area tracks numerosity. While these possibilities have entertained theoretical possibilities, they were limited in explaining empirical neural data.

More recently, two computational mechanisms with connections to neural data have gained traction. Park and Huber (Park & Huber, 2022) adopted the stimulus design originally developed by DeWind and colleagues (DeWind et al., 2015) where dot arrays vary in three independent dimensions: number, size, and spacing. Number refers to the number of dots. Size refers to the dimension that changes with the individual dot area while holding number constant, which automatically changes the total surface area of the all the dots (the area of each dot is designed to be identical within an array). Spacing refers to the dimension that changes with the placement of dots in an array; the dots can be placed close to each other (increasing density and decreasing field area), or they can be placed farther from each other (decreasing density and increasing field area). Park & Huber (Park & Huber, 2022) demonstrated that the summation of convolutional neuronal activities from center-surround cells across multiple-spatial scales minimizes the aggregate neural activity’s sensitivity to the spacing dimension and that divisive normalization between those center-surround neurons minimizes the aggregate neural activity’s sensitivity to the size dimension. A subsequent study suggested that this computational model makes unique predictions at the neural level and indeed demonstrated that the predictions are validated in an EEG study (Croteau et al., 2024). These findings suggest that *normalized local contrasts* of the image serve as the basis for numerosity perception.

In Paul and colleagues’ work (Paul et al., 2022), the authors used the population receptive field modeling (pRF) approach to study the representation of numerosity in the brain. Previously, the pRF approach has been used extensively to identify brain regions that show a tuned response to numerosities as well as to other magnitudes (Harvey et al., 2013, 2015). In this newer study, Paul and colleagues (Paul et al., 2022) modeled both tuned and monotonic responses to characterize how different brain regions may subserve different response functions, and they reported that activities in early visual areas are more aligned with monotonic response functions while parietal areas are more aligned with tuned functions. In their explanation for how numerosity encoding becomes invariant to other confounding magnitude dimensions, the authors identified that the summation of 2D Fourier power spectral density (across all orientations) up to the first harmonic closely tracks the numerosity of the image. In fact, their analysis showed that early visual cortical activities best match with this “aggregate Fourier power” more so than numerosity, from which the authors concludes that the sensory encoding of numerosity is based on human visual system’s approximation of a Fourier decomposition. It is worth noting that both accounts reject the idea that discrete numerical values are directly encoded in the visual system. According to this line of argument, numerical interpretation of these early sensory representations (e.g., normalized local contrast or Fourier power) is possibly enabled by a post-perceptual process (for a theoretical proposal on this, see Park, in press).

There are some reasons to doubt the aggregate Fourier account (Paul et al., 2022). First, it is unclear how the neural system would approximate a Fourier decomposition, given that Fourier decomposition relies on basis functions with infinite extent while biological systems are thought to rely on localized basis sets such as Difference-of-Gaussian or Gabor filters (see **Fig. 1**). Second, the computation of aggregate Fourier power requires the knowledge about the harmonic frequency, and how this can be achieved in the brain is questionable. It is unclear how the brain would figure that out and how their metric should be computed when there is no harmonic structure (e.g., in arrays composed of items without sharp edges). Third, some implementation choices in Paul and colleagues’ work are inconsistent with a typical practice in vision science. While the authors state that they computed Fourier power spectrum, their implementation indicates that amplitude instead of power was computed. In addition, while it is customary to average the power across orientations, the authors summed up the Fourier amplitude which could be a biased measure because more discrete Fourier coefficients lie at larger radial distance (i.e., higher frequencies). In addition to these theoretical and practical reasons, the current study raises empirical reasons to reject the aggregate Fourier account. If sensory encoding for numerosity is based on Fourier decomposition of the input image, early neural signals in response to FPE images should be entirely abolished. That was not the case. Although some disruption in the modulatory effect of magnitudes was observed at the prototypical C1 level, the visual-evoked potentials were robustly sensitive to the magnitudes (particularly *N*) throughout the time course across the posterior channels.

In defense of Paul and colleagues’ work (Paul et al., 2022), the aggregate Fourier account may simply be providing a mathematical lens for understanding spectral power analysis, rather than providing a biologically plausible model of numerosity encoding. After all, the biological system does not operate at the global level as Fourier decomposition does, and localized filters like the Difference-of-Gaussian or Gabor have long been acknowledged as ways to model visual receptive fields. For example, one can posit that the cortex does a Fourier-like decomposition of spectral power within a *localized* Gaussian window (Daugman, 1985). However, this view is still problematic for two reasons. First, it does not align well with the central claim concerning aggregate (band-limited) Fourier power in Paul and colleagues’ work (Paul et al., 2022). Therefore, the claim that aggregate Fourier power best aligns with numerosity will need a re-evaluation if such a spectral analysis done computed locally. Second, this localized Fourier-like decomposition would be conceptually identical to measuring energy captured by DoG or Gabor filters. However, (Park & Huber, 2022) demonstrated that energy from a layer of convolutional filters (without divisive normalization) is most correlated with the total surface area of the dot array, not numerosity. Only when divisive normalization was implemented, the energy encoded in the convolutional layer correlated with numerosity. Thus, the idea of Fourier decomposition in numerosity perception has limitations, even if it is to emphasize the notion of spectral analysis regardless of biological plausibility.

One noteworthy result from the EEG experiment was the overall polarity differences for the *N* effects between the VEPs for Intact and FPE images (**Fig. 4**). Observed waveforms are a result of the weighted sum of underlying activities from multiple neural generators that overlap over time. Therefore, it is difficult to determine if these differences between the patterns particularly in later latencies indicate differences in the later processing stages or is a result of differences in the earlier processing stages (Kappenman & Luck, 2012). Future studies using multiple neuroimaging techniques may be useful in identifying how the processing of Intact versus FPE images differ beyond early feedforward stages.

### Sensory representation of numerosity in coarse and fine visual channels

It is well-established that visual information is processed through two parallel retinogeniculate pathways (Livingstone & Hubel, 1988). The so-called M pathway starts from retinal ganglion cells that have large receptive fields which project to the magnocellular layer of the lateral geniculate nucleus (LGN). These cells are phasic and are sensitive to luminance contrast. In contrast, the P pathway starts from retinal ganglion cells that have small receptive fields which project to the parvocellular layer of the LGN. These cells tonic and are sensitive to chromatic contrast. The two pathways remain largely segregated in early cortical stages as well. In sum, the M pathway is a relatively fast visual channel that primarily engages low spatial frequency information, while the P pathway is a relatively slow visual channel that primarily engages high spatial frequency information.

Previous VEP studies show that manipulation of visual features that distinguish the two pathways modulate medial occipital activities, particularly the C1 component or a negative deflection around 100 ms (also referred to as the N1 in the visual-evoked potentials literature). Particularly, visual stimuli that favor the P pathway, such as isoluminant chromatic contrast with relatively high spatial frequency, is found to substantially modulate the C1 activity (Ellemberg et al., 2001; Foxe et al., 2008; Jones & Keck, 1978; Murray et al., 1987; Tobimatsu et al., 1995). In fact, when the stimuli maximally isolate the M pathway, using a low contrast luminance contrast in low spatial frequency, the prototypical C1 activity is abolished, while other modulatory effects remain observed such as earlier around 60-70 ms or later at the first positive deflection around 150 ms (Baseler & Sutter, 1997; Foxe et al., 2008; Hansen et al., 2011).

According to this rich literature concerning the contribution of P pathway to C1 activity, the FPE images disrupting C1 activity in the current study (see **Fig. 5**) suggests that the encoding of magnitude from the FPE images occurred mainly through the coarse spatial-scale information and not through the fine spatial-scale information. When this insight about the distinction between coarse and fine visual channels was implemented in the divisive normalization model, the simulation results were consistent with the neural data. Namely, the coarse channels, but not the fine channels, showed sensitivity to *N* in the FPE images (see **Fig. 7**). These findings suggest the possibility that sensory representation of numerosity is achieved through multiple visual channels.

### Predictive power of the divisive normalization model

One limitation of the current work is that, for simplicity, we simulated coarse and fine channels independently based on the spatial scale of the DoG filters. However, these channels in reality must work together to give rise to a coherent percept. How do the coarse and fine channels interact with each other to give rise to the mental representation of numerosity? Much empirical research has led to the idea that coarse channels engage with the stimulus earlier than the fine channels in the visual stream, which is referred to as coarse-to-fine processing (Hegdé, 2008; Hochstein & Ahissar, 2002). At the neural level, a subtype of V1 neurons responds exclusively for low spatial frequency with its firing rate peaking earlier than other subtype of neurons that responds exclusively for high spatial frequency (Skyberg et al., 2022). Studies investigating the dynamics of spatiotemporal receptive field characteristics also report that many V1 neurons are initially tuned broadly and that the tuning becomes sharper over the time course (Bredfeldt & Ringach, 2002; Frazor et al., 2004; Malone et al., 2007; Mazer et al., 2002; Purushothaman et al., 2014). At the behavioral level, previous studies tested the coarse-to-fine processing hypothesis by presenting to observers a sequence of low-pass to high-pass filtered images or a sequence of high-pass to low-pass filtered images in a categorization or identification task (Musel et al., 2014; Parker et al., 1992, 1997; Schyns & Oliva, 1994). Observers perform better when the coarse features of the image were presented prior to the fine features of the image. Consistent with these behavioral findings, human neuroscience studies show evidence of a rapid magnocellular projection to higher level brain areas, such as the prefrontal cortex (Bar, 2003; Kveraga et al., 2007; Petras et al., 2019; Peyrin et al., 2010). Based on these findings, one possible neurocomputational mechanism for coarse-to-fine processing may be gain modulation, in which one neuronal system modulates the sensitivity or gain of another neuronal system (Salinas & Sejnowski, 2001). According to this idea, our model could be modified so that the coarse channels (or the “blob” layers in **Fig. 7**), which engages the visual input faster than the fine channels (the “edge” layers), modulate the activity of fine channels for the sensory representation of the stimulus. Of course, a more complex computational mechanisms in a deep neural network frame may be entertained as well (Zou et al., 2023). Future work should explore these possibilities.

Despite the simplistic modeling of coarse and fine channels, our divisive normalization model made one clear, novel prediction. If the processing of fine channel is disrupted, the fidelity of sensory representation will be impacted by this disruption for Intact images but not for FPE images. Indeed, backward masking which is known to disrupt fine processing and preserve coarse processing worsened the numerical acuity only for Intact images (**Fig. 8**). In a previous study, divisive normalization model of numerosity perception made a novel prediction about how the neural basis of coherence illusion may differ based on the source of coherence, and this novel prediction was validated with neural data (Croteau et al., 2024). The current study showcases another important empirical validation of a novel prediction generated by the computational model.

## Conclusion

We conducted an integrated neural, behavioral, and computational investigation on the sensory representation of numerosity. Our results reject the hypothesis that numerosity representation is based on Fourier spectral power; the findings are rather consistent with the idea that numerosity representation is achieved through normalized local contrast. Moreover, the current results demonstrate the importance of considering visual channels across multiple spatial scales to gain a comprehensive understanding of numerosity perception and indicate a coarse-to-fine processing in perception of numerosity. In sum, this study advances our understanding of the neural mechanism for representational invariance of numerosity.

## Acknowledgements

The authors thank Rachael McCollum, Emily Isaacs, Matthew Gillespie, Nathalie Valentin, Marina Morcos, Victoria Schille, Zachary Parsons, Hong Mai Ngo, Kirthi Sivakumar for data collection and Larry Liu for technical assistance. We thank Dr. David Huber, Dr. Hansem Sohn for invaluable comments on the paper.

## Data availability

The MATLAB code for the divisive normalization model of numerosity perception can be found in the following public repository: https://osf.io/4rwjs/.

## Author contributions

Joonkoo Park: Conceptualization, Methodology, Software, Formal analysis, Writing. Shimin Hu: Formal analysis, Writing.

